# Lysergic Acid Diethylamide Alters the Effects of Brain Stimulation in Rodents

**DOI:** 10.1101/2022.10.31.514588

**Authors:** Lucas Dwiel, Angela Henricks, Elise Bragg, Jeff Nicol, Jiang Gui, Wilder Doucette

**Affiliations:** Department of Psychiatry, Geisel School of Medicine, Dartmouth College, Hanover, NH; Department of Psychology, Washington State University, Pullman, WA; Department of Psychiatry, Dartmouth-Hitchcock Medical Center, Lebanon, NH; Geisel School of Medicine, Dartmouth College, Hanover, NH; Department of Biomedical and Data Sciences, Geisel School of Medicine, Dartmouth College, Hanover, NH

**Keywords:** psychedelics, neuromodulation, electrophysiology, rodent

## Abstract

**Background:** Psychedelic drugs have resurged in neuroscience and psychiatry with promising success in psychedelic-assisted therapy for the treatment of anxiety, depression, and addiction. At the cellular level, psychedelic drugs elicit neuroplastic processes 24 hours after administration, priming neural circuits for change. The acute effects of psychedelic drugs are well characterized with functional imaging and neural oscillations showing an increase in the entropy of spontaneous cortical activity.

**Hypotheses:** We hypothesized that cortical-striatal oscillations recorded in rats would confirm the effects of psychedelic drugs. We also hypothesized that brain stimulation delivered 24 hours after LSD administration would lead to different effects than brain stimulation alone.

**Methods:** We recorded local field potential (LFP) oscillations from rats following lysergic acid diethylamide (LSD) or saline administration and determined if exposure to these treatments altered the effect of a targeted intervention (brain stimulation) 24 hours later.

**Results:** We confirmed acutely decreased low frequency power across the brain when rats are given LSD. We also demonstrated these altered states return to baseline after 24 hours. Brain stimulation applied in the previously reported window of heightened neuroplasticity produced distinct shifts in brain state compared to brain stimulation applied 24 hours after saline.

**Conclusions:** Despite the acute effects of LSD disappearing after 24 hours, there are still latent effects that interact with brain stimulation to create larger and distinct changes in brain activity compared to brain stimulation alone. Our proof-of-concept findings are the first to suggest that psychedelic drugs could work in combination with brain stimulation to achieve enhanced effects on brain activity and future work will assess impacts on stimulation induced changes in behavior.

## Introduction

Psychedelic drugs have achieved breakthrough status from the Food and Drug Administration after remarkable success for the treatment of depression^1^ and post-traumatic stress disorder.^2^ Research with classic psychedelic drugs, like psilocybin and lysergic acid diethylamide (LSD), has demonstrated a correlation between acute subjective effects (e.g., a mystical experience) and reported therapeutic effects.^3, 4^ However, it remains unclear if the mystical experience or other subjective effects are merely a proxy of achieving a therapeutic dose related to another mechanism or if the experience itself plays a necessary role in therapeutic efficacy.^4^ The brain activity changes that coincide with the acute subjective effects have been described in human and animal studies using functional brain imaging and electrophysiologic approaches. These systems-level brain activity readouts indicate that, acutely, psychedelic drugs disrupt the coordination of brain activity within and between brain regions. For example, imaging studies in humans reveal reduced connectivity between nodes of the default mode network ^5, 6^ and recordings of neural oscillations (e.g,. electroencephalography, EEG; or local field potentials, LFP) show that psychedelic drugs reduce power across frequencies, particularly low frequencies^7^. Beyond the subjective effects and immediate changes in brain activity induced by psychedelic drugs, preclinical work has uncovered alternative, or complementary, mechanisms of enhanced neural plasticity.

Both *in vitro* and *in vivo* psychedelic drugs have been shown to open a window of enhanced neural plasticity^8^ as well as produce anti-inflammatory effects^9–11^. Both mechanisms represent biologically plausible mechanisms of therapeutic efficacy in humans. The plastogenic properties of psychedelic drugs include the growth of new neuronal processes, like dendritic spines, with about one third of these spines becoming persistent, functional synapses.^12^ It is theorized that the enhanced neural plasticity induced by psychedelic drugs could contribute to the reported persistent therapeutic effects in patients. For example, in a pre-clinical rat model ketamine acutely ameliorates depressive behaviors before inducing new spine growth, with the long-term effects of ketamine dependent on the newly formed spines.^13^ This work suggests that although there are potentially clinically meaningful acute effects of psychedelic drugs, the long term success of psychedelic drugs to change behavior is primarily mediated by the growth of new functional connections between neurons.

We theorize that directed interventions such as psychotherapy or brain stimulation may be able to direct the neuroplastic changes induced by psychedelic drugs to create lasting changes in synaptic organization and ultimately in behavior. We hypothesized that cortical-striatal oscillations recorded in rats would confirm the well characterized effects of psychedelic drugs (tested here with LSD) and show for the first time the systems-level brain activity changes corresponding to the period of enhanced neural plasticity approximately 24 hours after LSD administration. Next, we characterized how brain stimulation targeting the rat infralimbic cortex (a medial frontal region) altered cortical-striatal oscillations when rats were given LSD or saline 24 hours before stimulation. This allowed us to determine if LSD allows brain stimulation to have different and/or larger effects on brain activity than when saline was given before stimulation.

We used a rodent model to remove the unavoidable biases inherent to psychedelic research in humans, and used brain stimulation to manipulate brain activity rather than psychotherapy, to begin probing the potential interaction of an external manipulation of brain activity and the window of psychedelic induced neural plasticity. The results presented below begin to reveal the potential interaction between an external manipulation of brain activity and the window of enhanced neural plasticity created by psychedelic drugs. Beyond probing the systems-level brain activity mechanisms that could underlie the clinical combination of psychedelic drugs and psychotherapy, this work also investigates the potential interaction of psychedelic drugs and brain stimulation—a potential new therapeutic approach.

## Methods

### Drugs

We mixed LSD with sterile saline and delivered the drug via an intraperitoneal injection at 0.15 mg/kg. We chose this dose to be large enough to produce changes in behavior^14^ and early gene expression^15^.

### Cohorts

We used two cohorts of Sprague Dawley rats (Charles River) of each sex for these experiments. The first cohort (n = 11; 5 male and 6 female) came from the control group of a separate experiment. The second cohort (n = 5; 2 male and 3 female) had been trained on a delayed discounting task for a separate experiment.^16^ Both cohorts were given at least one month before incorporating into these experiments. All animals arrived at 60 days old and were approximately 120-180 days old at the start of these experiments. All experiments were carried out in accordance with the National Institute of Health Guide for the Care and Use of Laboratory Animals (NIH Publications No. 80–23) and were approved by the Institutional Animal Care and Use Committee of Dartmouth College.

### Electrode implantation

After habituating the rats to the animal facility for 1 week, we anesthetized them with isoflourane/oxygen (5% isoflourane for induction and 2% for maintenance) and stereotactically implanted them with one of two custom electrode arrays to record LFPs and deliver electrical stimulation. Briefly, these electrodes use a custom milled plastic base with polyimide tubing that guides two wires to each brain location (one nichrome wire for recording, 50 μm, and the other stainless steel wire for stimulating, 125 μm). We implanted the electrodes such that the polyimide tubing and wires were below the skull, with the plastic base attached to the skull surface with acrylic cement. The first array for the first cohort targeted the bilateral infralimbic cortex—IL (AP 3.4 mm; ML ±0.75 mm; and DV -5.0 mm), and nucleus accumbens shell—NAcS (AP 1.2 mm; ML ±1.0; and DV -7.6 mm). The second array for the second cohort targeted the same bilateral IL and NAcS as well as orbitofrontal cortex—OFC (AP 3.4 mm; ML ±3.0 mm; and DV -6.0 mm), and the nucleus accumbens core—NAcC (AP 1.2 mm; ML ±2.4 mm; and DV -7.6 mm). All coordinates are relative to bregma.

### Local field potential processing

We assessed the brain states of the rats using LFP features, power and imaginary coherence, within 6 frequency ranges: delta (Δ) = 1-4 Hz, theta (θ) = 5-10 Hz, alpha (α) = 11-14 Hz, beta (β) = 15-30 Hz, low gamma (lγ) = 45-65 Hz, and high gamma (hγ) = 70-90 Hz. We used the imaginary component of coherence to minimize the influence of volume conduction and a common reference.^17^ From the 4 recording locations (bilateral IL and NAcS) in the first cohort, we obtained 24 power features and 36 coherence features, and from the 8 recording locations (bilateral IL, OFC, NAcC, and NAcS) in the second cohort, we obtained 48 power features and 168 coherence features, providing a total of 60 features from 4 recording sites or 216 features from 8 recording sites to describe brain states. Apart from the use of imaginary coherence, all signal processing was done as previously described.^18^ All recordings were done with the rats tethered in operant boxes (MedAssociates, Fairfax, Vermont) with a house light on for video recording.

### Measuring the acute effects of LSD

In the first cohort of rats we recorded LFPs from bilateral NAcS and IL for 90 minutes during free behavior on two separate days to serve as baseline data. We then recorded intervention sessions in which we recorded 10 minutes to obtain baseline brain activity, administered LSD (n = 6; 3 male and 3 female) or saline (n = 5; 2 male and 3 female), and then recorded LFPs for another 90 minutes (**Supplemental Figure 2**). To quantify how an injection of LSD or saline (SAL) changed brain activity (**Supplemental Figure 3**) we subtracted baseline features (averaged from the two baseline recordings and the 10-minute, pre-injection baseline) from every 3-second feature quantification from the post-injection period. We then trained logistic regressions on each combination of 5 LSD and 4 SAL rats, leaving out one rat from each group for testing (leave-one out, LOO), resulting in 30 iterations of model building and testing. We used the distribution of these model performances to estimate the magnitude of the difference between brain activity from LSD or SAL exposed rats. We then built models using each LFP feature separately to assess individual features. We repeated the model building using data collected from the same rats 24 hours after injection to determine if the effects of LSD persist. As LSD causes acute ataxia, we limited the data used for modeling to the last 30 minutes of the recording when both groups of rats had similar levels of resting behavior (**Supplemental Figure 1**) manually scored with resting defined as intervals in which the rats were either asleep or awake and not moving (e.g., not locomoting, grooming, and rearing).

### Measuring the effects of brain stimulation 24 hours after LSD

In the second cohort of rats, we first administered SAL and waited 24 hours. We then recorded LFPs from bilateral NAcS, NAcC, IL, and OFC for 10 minutes to obtain baseline brain activity and administered IL stimulation (monopolar, 130 Hz, 90 μs pulse width, and 200 μA) for 120 minutes while recording (**Supplemental Figure 4** and **5**). We repeated this pairing of SAL pretreatment and 120 minutes of IL stimulation once a day for three days. After these SAL+stim sessions we allowed at least 1 week for stimulation effects to wash out before injecting LSD (0.15 mg/kg i.p.), waiting 24 hours, and repeating the record/stimulation paradigm once (**Supplemental Figure 4** and **6**). First, we trained logistic regressions to differentiate between brain activity at baseline and during stimulation for both SAL and LSD (i.e., differentiating brain activity from the 10 minutes before stimulation and the 120 minutes during stimulation). Since we had multiple sessions of SAL+stim per rat, we downsampled the data used from each rat to match the sample sizes from the single LSD+stim sessions. Next, we built logistic regressions to differentiate directly between the effects of brain stimulation with LSD or SAL given 24 hours earlier (i.e., LSD+stim vs. SAL+stim) by subtracting the average baseline brain activity from 3-second samples of brain activity during stimulation (**Supplemental Figures 7-8**).

### Model performance and significance

To assess the performance of these models, we used the receiver operator characteristic curve obtained by plotting the true positive rate (TPR) vs. the false positive rate (FPR) and then finding the area under this curve (area under the receiver operator characteristic curve, AUROC) such that an AUROC of 1 would be perfect performance and 0.5 would be the by-chance performance. The AUROC therefore reflects the difference in mean and standard deviation of all of the features across groups (**Supplemental Figure 3**). To estimate by-chance model performance we also repeated the model building and testing on permuted data in which the assignment of brain data to a group (i.e., LSD or SAL) was randomized. We can then compare the mean model performance to the distribution of permuted model performances and calculate a p-value.^19^ For all model building we used 80% of the data, with equal contributions from each rat, to train the models and tested them on the remaining 20%.

## Results

### LSD leads to acute decreases in power and mixed effects in coherence

LSD leads to acute changes in LFPs that are differentiable from the changes induced by SAL using logistic regressions with LOO testing (acute = 0.79±0.02 vs. permuted = 0.5±0.04; **Fig 1A**). However, these changes in LFPs disappear within 24 hours (24 hrs = 0.45±0.037; **Fig 1C**). Building logistic regression models with one LFP feature at a time revealed that LSD leads to acute decreases in NAcS and IL power across all frequency ranges and a mixture of increased and decreased coherence (**Fig 1B**). For a more detailed visualization of the difference between the effects of LSD and SAL on power and coherence, see **Supplemental Fig 2**. The average performance of each feature is also listed in **Table 1**.

**Figure 1.**
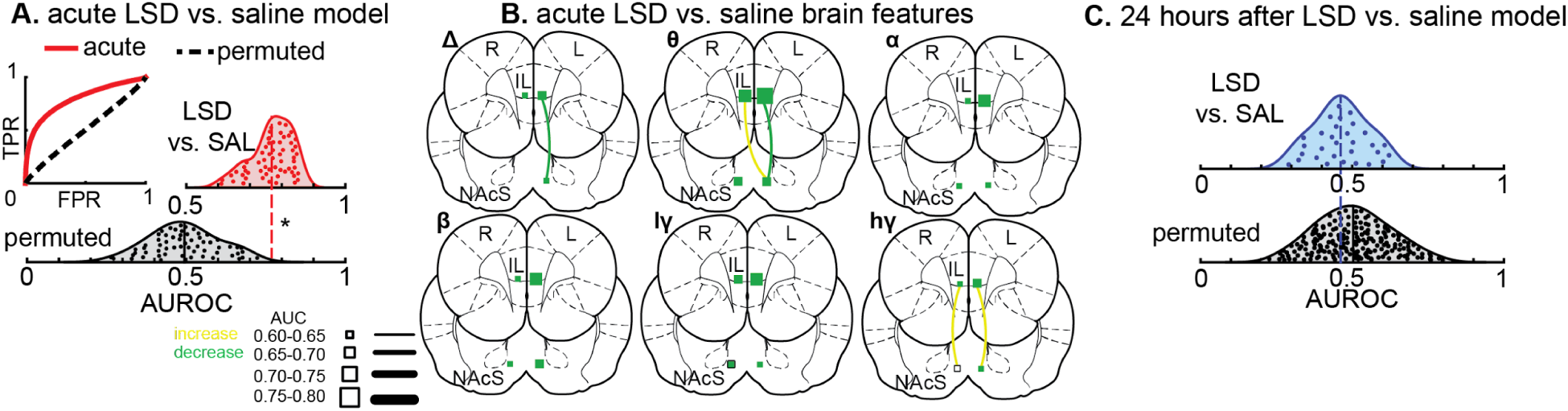
Compared to saline, LSD leads to global decreases in power and mixed changes in coherence; however, these differences disappear within 24 hours. **A** Rats given LSD can be differentiated from those given saline (SAL) using all of the acute changes in power and coherence between SAL and LSD in logistic regressions (acute; red solid line) compared to permuted data (black dashed line). True positive rate, TPR; false positive rate, FPR.**B** The performance of logistic regressions built using individual brain activity features with squares representing power at a specific brain location and lines connecting brain regions representing coherence between specific brain regions. The size of the square or weight of the line indicates the model performance (area under the receiver operator characteristic curve; AUROC). N.B.: all coherence feature performances fall within the same range (0.60-0.65). Green indicates a relative decrease in that brain feature with LSD compared to SAL and yellow indicates a relative increase. The data summarized here are shown in more detail in Supplemental Figure 2. Right, R; left, L; infralimbic cortex, IL; nucleus accumbens shell, NAcS; delta, Δ; theta, θ; alpha, α; beta, β; low gamma, lγ; high gamma, hγ; true positive rate, TPR; false positive rate, FPR. **C** Rats given LSD are indistinguishable from rats given SAL 24 hours after drug administration (blue distribution) as compared to permuted data (black distribution). * p < 0.05

**Figure 2.**
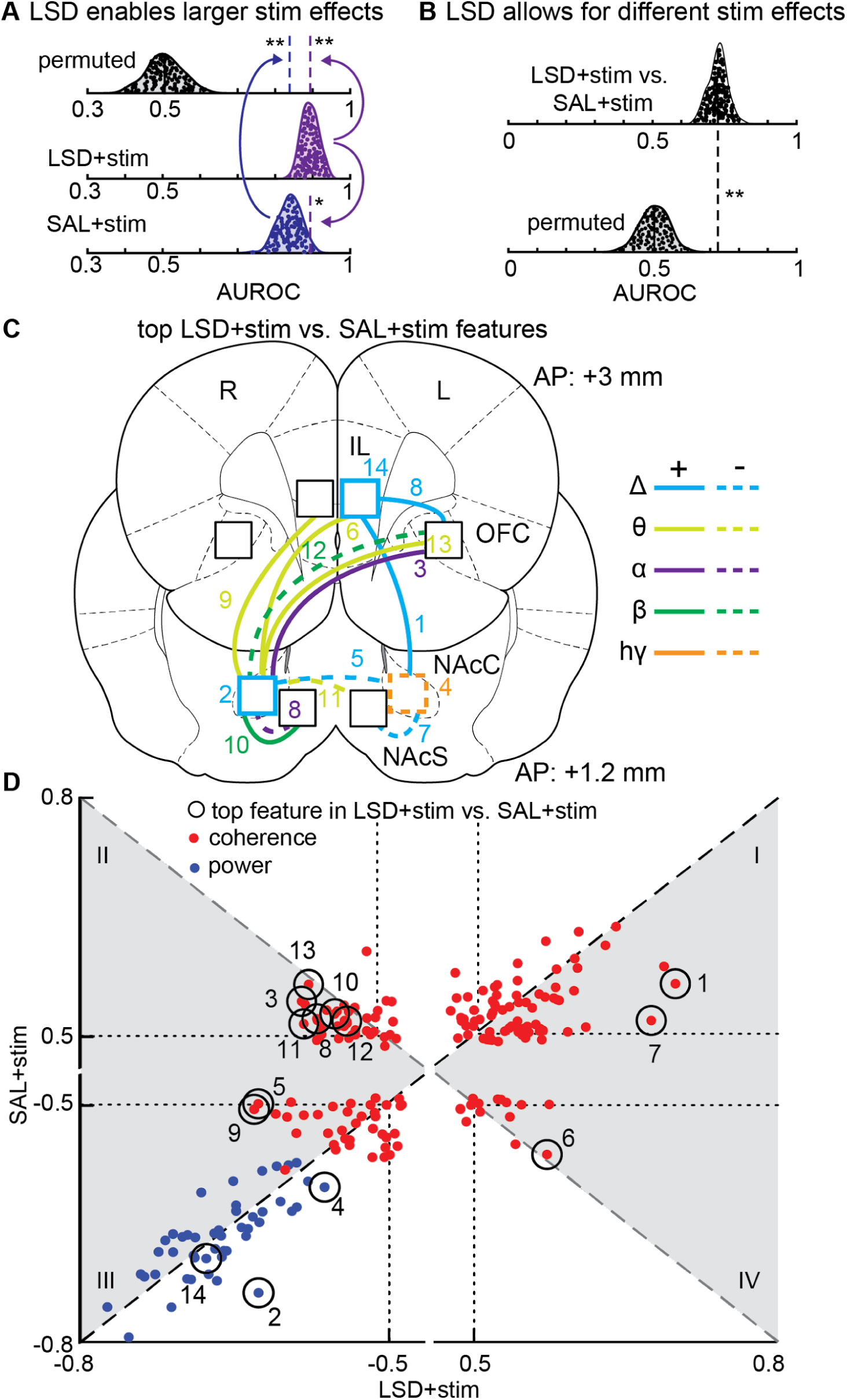
LSD interacts with brain stimulation to create larger and different changes in brain activity. **A** Logistic regressions built to distinguish between baseline brain activity and stimulated brain activity perform better when the rats were given LSD 24 hours before stimulation (LSD+stim = 0.90±0.005; gray solid line) compared to saline (SAL+stim = 0.84±0.006; solid black line). Both LSD+stim and SAL+stim out-performed permuted data (0.49±0.01; dashed black line). All model performances are reported as the mean±95% confidence interval of the distributions of the area under the receiver operator characteristic curves (AUROCs). **B** Logistic regressions built to distinguish the change in brain activity seen in LSD+stim vs. SAL+stim (0.90±0.004; solid line) out-perform permuted data (0.50±0.01; dashed line). All model performances are reported as the mean±95% confidence interval of the distributions of AUROCs. **C** Visualization of the top features from distinguishing LSD+stim from SAL+stim (**B**) superimposed on the rat brain. Brain features came from left (L) and right (R) infralimbic cortex (IL), orbitofrontal cortex (OFC), and nucleus accumbens core (NAcC) and shell (NAcS). Outlines represent power at a given brain region and lines between brain regions represent coherence. Solid lines/squares indicate a relative increase in that brain feature with LSD+stim compared to SAL+stim and dashed lines/squares indicate a relative decrease. The color of the line/outline indicates the frequency range. **D** Visualization of all single feature model performances from distinguishing LSD+stim (x-axis) or SAL+stim (y-axis) from baseline (**A**) with positive values indicating a relative increase in that feature from baseline and negative values indicating a relative decrease in that feature from baseline. Features are numbered in descending order from the highest |AUROC| (**Table 2**). Blue dots are power features and red dots are coherence features. Circled features were the top performers in the models distinguishing between LSD+stim and SAL+stim (**B**). Light gray background highlights features with a higher |AUROC| in LSD+stim than SAL+stim; white background highlights features with lower |AUROC| in LSD+stim models than SAL+stim models. Features that fall along the dashed gray diagonal line have opposite correlations in LSD+stim compared to SAL+stim and features that fall along the dashed black diagonal line have correlations in the same direction in LSD+stim models compared to SAL+stim models. Dotted black lines delineate the 4 quadrants (I, II, III, and IV) of |AUROC|>0.5 for both models. * p < 0.05, ** p < 0.01

**Table 1.**
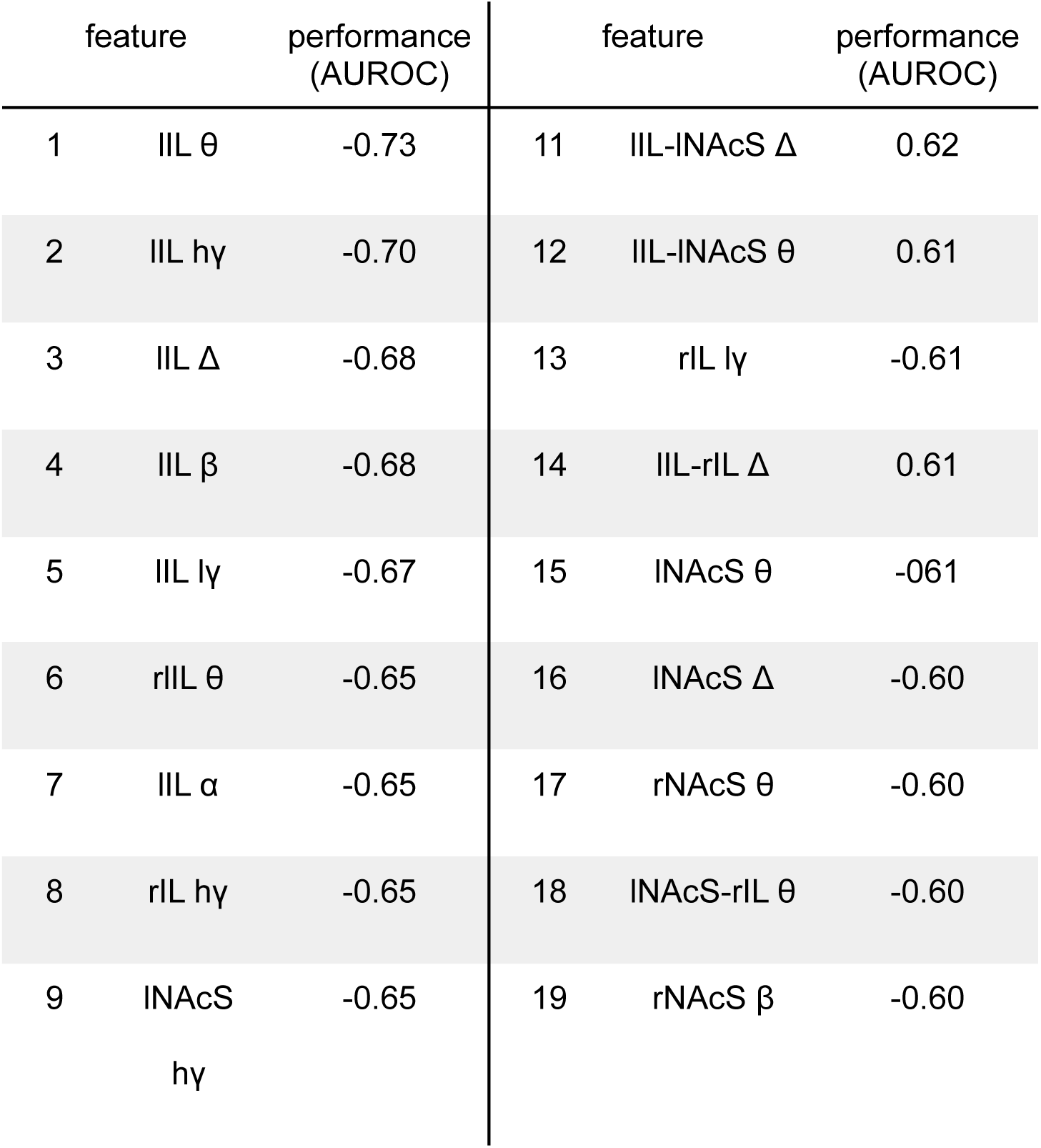
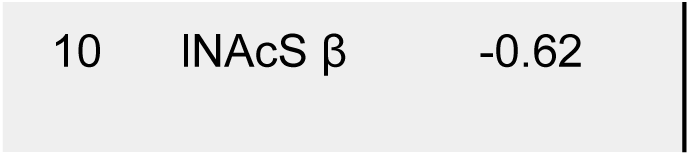
Brain features most predictive of whether a rat was given LSD vs. saline. Features are notated with a lowercase letter indicating hemisphere (left, l; right, r). A single brain region indicates power from that region while two regions combined with a hyphen indicates coherence between those regions. For example, the first feature (lIL θ) is theta power from left infralimbic cortex. NAcC, nucleus accumbens core; NAcS, nucleus accumbens shell; IL, infralimbic cortex; OFC, orbitofrontal cortex; Δ, delta; θ, theta; α, alpha; β, beta; lγ, low gamma; hγ, high gamma.

| feature performance<br>(AUROC) |  |  | feature performance<br>(AUROC) |  |  |
| --- | --- | --- | --- | --- | --- |
| 1 | IIL $\theta$ | -0.73 | 11 | IIL-INAcS $\Delta$ | 0.62 |
| 2 | IIL hy | -0.70 | 12 | IIL-INAcS $\theta$ | 0.61 |
| 3 | IIL $\Delta$ | -0.68 | 13 | rIL ly | -0.61 |
| 4 | IIL $\beta$ | -0.68 | 14 | IIL-rIL $\Delta$ | 0.61 |
| 5 | IIL ly | -0.67 | 15 | INAcS $\theta$ | -0.61 |
| 6 | rIL $\theta$ | -0.65 | 16 | INAcS $\Delta$ | -0.60 |
| 7 | IIL $\alpha$ | -0.65 | 17 | rNAcS $\theta$ | -0.60 |
| 8 | rIL hy | -0.65 | 18 | INAcS-rIL $\theta$ | -0.60 |
| 9 | INAcS<br>hy | -0.65 | 19 | rNAcS $\beta$ | -0.60 |

|  |  |  |
| --- | --- | --- |
| 10 | INAcS $\beta$ | -0.62 |

**Table 2.** Top performing features in distinguishing the effects of brain stimulation with LSD given 24 hours previously from the effects of brain stimulation with saline (SAL) given 24 hours previously (LSD+stim vs. SAL+stim). For comparison, the performance of each of these features in the two models comparing intervention to baseline (LSD+stim and SAL+stim) are displayed to the right. All performances are reported as the area under the receiver operator characteristic curve (AUROC) with the sign indicating the direction of the correlation. Features are notated with a lowercase letter indicating hemisphere (left, l; right, r). A single brain region indicates power from that region while two regions combined with a hyphen indicates coherence between those regions. For example, the third best feature (lOFC-rNAcC α) is alpha coherence between left orbitofrontal cortex and right nucleus accumbens core; the negative sign indicates that higher coherence correlates with SAL+stim versus LSD+stim (i.e., LSD+stim leads to lower lOFC-rNAcC α coherence than SAL+stim). NAcC, nucleus accumbens core; NAcS, nucleus accumbens shell; IL, infralimbic cortex; OFC, orbitofrontal cortex; Δ, delta; θ, theta; α, alpha; β, beta; hγ, high gamma.

|  |  | LSD+stim |  |  |
| --- | --- | --- | --- | --- |
|  |  | vs. |  |  |
| feature |  | SAL+stim | SAL+stim | LSD+stim |
|  |  | (AUROC) | (AUROC) | (AUROC) |
| 1 | lIL-INAcC $\Delta$ | 0.66 | 0.57 | 0.70 |
| 2 | rNAcC $\Delta$ | 0.63 | -0.75 | -0.63 |
| 3 | lOFC-rNAcC $\alpha$ | 0.63 | 0.55 | -0.58 |
| 4 | INAcC hγ | -0.62 | -0.61 | -0.56 |
| 5 | INAcC-rNAcC $\Delta$ | -0.62 | -0.5 | -0.63 |
| 6 | lIL-rNAcC $\theta$ | 0.62 | -0.57 | 0.57 |
| 7 | INAcS-INAcC $\Delta$ | -0.62 | 0.52 | 0.68 |
| 8 | rNAcS-rNAcC $\alpha$ | -0.62 | 0.52 | -0.56 |
| 9 | rIL-rNAcC $\theta$ | 0.61 | 0.51 | -0.64 |
| 10 | rNAcS-rNAcC $\beta$ | 0.61 | 0.53 | -0.54 |
| 11 | INAcS-rNAcC $\theta$ | -0.61 | 0.52 | -0.58 |
| 12 | IOFC-rNAcC $\beta$ | -0.61 | 0.52 | -0.53 |
| 13 | IOFC-rNAcC $\theta$ | -0.61 | 0.57 | -0.57 |
| 14 | IIL $\Delta$ | 0.60 | -0.70 | -0.68 |

### LSD synergizes with brain stimulation

Despite brain activity being indistinguishable between rats given LSD or SAL 24 hours prior (**Fig 1C**), IL stimulation is able to create larger changes in brain activity from baseline in rats given LSD (0.90±0.005; **Fig 2A**) compared to SAL (0.84±0.006; **Fig 2A**). Further, in the rats given LSD 24 hour before stimulation, the effects of stimulation are distinguishable from the effects of stimulation in rats given SAL (0.73±0.006; **Fig 2B**). Visualizing the performance of each feature in distinguishing LSD+stim vs. SAL+stim highlights that giving a rat LSD 24 hours before brain stimulation primarily leads to differential effects on coherence with increased delta cortico-striatal coherence, decreased delta intra-striatal coherence, increases in theta and alpha cortico-striatal coherence, and selective delta power increases and a high gamma power decrease. (**Fig 2C** and **Table 2**).

Plotting the performance of each feature in differentiating either LSD+stim or SAL+stim from baseline brain activity highlights that regardless of whether a rat was given LSD or SAL, IL stimulation led to broad decreases in power (blue dots; **Fig 2D**) and mixed effects in coherence (red dots; **Fig 2D**). This manifests as all power features falling along the black dashed line in quadrant III and coherence features existing in all quadrants. The features that fall along the black dashed line (quadrants I and III) change in the same direction relative to baseline activity, although to different degrees. For example, the second best performing single feature (Δ power at the right NAcC, AUC = -0.63; **Table 2**) had large decreases in rats given SAL before stimulation and modest decreases in rats given LSD before stimulation (-0.75 vs. -0.63; **Table 2**). The features along the gray dashed line (quadrants II and IV) change in opposite directions relative to baseline activity. For example, the third best performing feature (α coherence between left OFC and right NAcC, AUC = 0.64; **Table 2**) increased slightly when the rat was given SAL 24 hours prior, and decreased slightly when given LSD 24 hours prior (0.55 vs. -0.58; **Table 2**).

## Discussion

These results further characterize the acute changes induced by psychedelic drugs (e.g., LSD) on cortical-striatal brain states and demonstrate these altered states return to baseline after 24 hours. However, a focal intervention applied in the reported window of heightened neuroplasticity (24 hours after LSD) produced distinct shifts in brain state compared to brain stimulation applied 24 hours after SAL. These results suggest that despite the neuroplastic changes reported 24 hours after a single dose of LSD,^8^ we were unable to identify any systems-level brain activity differences using neural oscillations. However, when we applied brain stimulation, we found significant differences in brain activity changes depending on whether the rat was given LSD or SAL 24 hours prior. Thus, we theorize that the LSD-induced neuroplastic changes led to a latent state with an altered potential to respond to external manipulations or interventions (**Figure 3**). We believe that the mechanisms behind the interaction of psychedelic drugs and psychotherapy are likely also at play when LSD is combined with brain stimulation. We acknowledge that brain stimulation does not model the effects of psychotherapy directly, but rather, serves as an alternative method to externally direct brain activity changes in rodents.

**Figure 3.**
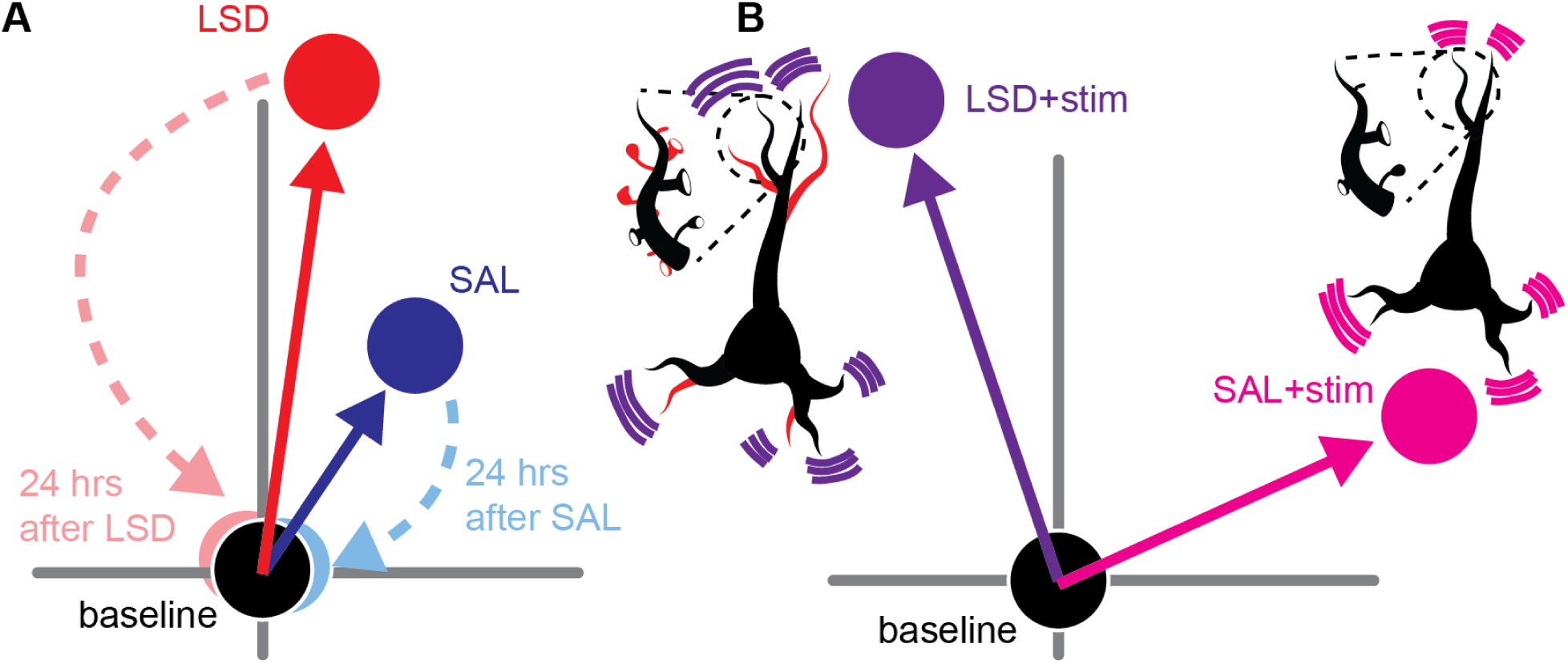
Conceptual diagram of the acute effects of LSD and SAL and the interaction between LSD and brain stimulation. **A** Acute LSD and SAL lead to differentiable effects in brain activity from baseline represented by the distance between the circles representing the brain state space during baseline (black circle) and after LSD injection (red circle and arrow) or after SAL injection (blue circle and arrow). 24 hours after injections, brain activity is no longer differentiable between groups (24 hours after LSD, light red circle and dashed arrow; and 24 hours after SAL, light blue circle and dashed arrow). **B** Brain stimulation applied 24 hours after LSD or SAL injections leads to large and different changes in brain activity. Brain stimulation after LSD (purple circle and arrow) may be interacting with the induced neural plasticity (red dendrites and spines in inset neuron) leading to different changes in brain activity than stimulation after SAL (pink circle and arrow).LSD+stim has been drawn in a different quadrant than SAL+stim to illustrate that LSD+stim leads to different changes in brain features, including changing brain features in opposite directions (Figure 2D and **Table 2**).

### Acute electrophysiological effects of LSD

Although there is a paucity of studies examining the acute effects of psychedelic drugs on intracranial LFPs, our findings of broad decreases in power and mixed disruptions in coherence (**Fig 1**), particularly at low frequencies, align with the reported decrease in accumbal high gamma power^20^ and the cortical effects of LSD (measured by epidural electrodes)^4, 21^ found in previous studies. There is also evidence of the same broad decreases in low frequency power from rodents given other serotonergic psychedelic drugs, like psilocybin/psilocin,^21^ 1-(2,5-dimethoxy-4-iodophenyl-2-aminopropane) (DOI),^22, 23^ and 5-Methoxy-N,N-dimethyltryptamine (5-MeO-DMT)^24^ although the effects of 5-MeO-DMT could not be replicated in freely behaving, non-anesthetized, rats^25^.

Magnetoencephalography (MEG) in humans given LSD has revealed a similar signature of broadband decreases in power across the brain with the largest effects occurring at lower frequencies paired with mixed effects on connectivity.^6, 26–29^ Similarly, the acute effects of LSD have been measured in humans using functional magnetic resonance imaging^27, 30–32^ and electroencephalography,^33, 34^ again finding disruptions in low frequency activity and widespread changes in connectivity. It is difficult to translate these findings to specific alterations in LFPs^35–37;^ however, there is convergence to an increase of disorder in the brain through the serotonin (5HT) 2A receptor.^38, 39^

The broadband decrease in power, particularly at lower frequencies, has been correlated to the subjective effects of LSD^27^ and there is also evidence that decreases in low frequency power is indicative of increased neural activity^40^. Similarly, the acute disruption of functional connectivity measured with functional magnetic resonance imaging (fMRI) also correlates with the subjective experience of LSD.^31, 41^ Further, the degree of entropy induced by LSD is predictive of long-term personality changes.^42^ Despite being predictive of future personality change, it could be the case that the acute changes in brain activity are a proxy measure of an adequate dose required to induce personality change rather than a cause of enhanced plasticity. There is evidence that the acute changes in subjective experience and brain activity as well as the the long-term enhanced neuroplasticity are mediated by the 5HT-2A receptor such that antagonists like ketanserin are able to significantly attenuate if not ablate these effects.^43–46^ Although the subjective experience, the changes in brain activity, and the enhanced neuroplasticity are related to one-another through their dependence on 5HT-2A receptor activation, whether any of these effects of psychedelic drugs are causally related or parallel processes remains to be elucidated.

### Interaction between LSD and brain stimulation

To our knowledge, there have been no publications pairing brain stimulation with LSD in freely behaving animals. However, there is a growing pool of evidence that ketamine is able to facilitate long term potentiation both *in vitro*^47–49^ and *in vivo*^50, 51^, with the discrepancies between studies being attributed to differing doses of ketamine, differing time points, the concurrent use of anesthetics, and the intrinsic differences between *in vitro* and *in vivo* electrophysiology.

Although there are non-trivial differences between ketamine and classical psychedelic drugs like LSD, there is also emerging evidence these drugs activate the same downstream molecular pathways regulating neuroplasticity.^52^ LSD has been paired with intracranial self stimulation (ICSS) in both rats ^53^ and mice ^54^, and in both cases LSD had no effect on the ICSS either acutely or after the acute effects wore off (100 minutes in rats and 24 hours in mice after LSD administration). This suggests that even if pretreatment with LSD allows brain stimulation to have different effects on brain activity, it is not changing the behavioral reinforcing effects of brain stimulation targeted to the brain reward pathway.

In particular, classical psychedelic drugs like LSD and psilocybin are known to induce rapid and persistent growth of dendritic spines.^8, 12^ The exact mechanism behind this plastogenesis is still an area of rich research, but it is likely that it is at least in part mediated by 5HT_2A_R, brain-derived neurotrophic factor (BDNF), tyrosine receptor kinase B (trkB), and mammalian target of rapamyacin signaling pathways (mTOR)^44, 55^ [for reviews:^56, 57^] as well as long term changes in gene expression^58^.

Although we found that LSD+stim led to larger changes in brain activity compared to SAL+stim (**Fig 2A**) and the LSD+stim changes were easily distinguishable from SAL+stim (**Fig 2B**), about half of the features changed in the same direction, albeit with differences in magnitude, between LSD+stim and SAL+stim. The other half of the features changed in opposite directions (e.g., decreased with LSD+stim and increased with SAL+stim), but these features had AUROCs <0.60 when distinguishing from baseline brain activity. Therefore, it appears that LSD not only modulates the effects of brain stimulation but also has the potential to induce opposite and novel changes in brain activity compared to brain stimulation alone.

While the presented experiments were not designed to determine if LSD lengthens the duration of the effects of brain stimulation, it is important for this to be evaluated in future studies. It is also important to determine if the LSD+stim induced changes in brain activity have a behavioral correlate. Currently, brain stimulation is either delivered chronically in the case of invasive deep brain stimulation or over long courses of non-invasive (e.g., transcranial magnetic stimulation; TMS) intervention sessions .^59–61^ This approach could shorten treatment courses or increase response rates through facilitation of changes in structural connectivity and/or synaptic weights. A similar combinatorial approach using TMSpharmacologic augmentation of synaptic plasticity with D-cylcoserine (a partial N-methyl-D-aspartate agonist) recently demonstrated increased response rates in patients with depression.^62^ Pairing plastogenic agents with brain stimulation may also help non-invasive stimulation overcome the high relapse rates currently limiting these interventions from making a larger clinical impact.

### Limitations and future directions

Although this work provides a proof-of-concept that LSD interacts with brain stimulation, it must be noted that we do not know by what mechanisms this interaction is occurring. Our intention for giving stimulation 24 hours after LSD administration was to allow brain stimulation to interact with the molecular mechanisms underpinning the neuroplastic effects of LSD while they are still active. However, at 24 hours, despite no detectable LSD in plasma, no behavioral changes, and no differences in brain activity, there may still be on-going acute effects of LSD due to its binding persistence^63^ that could be contributing to the interaction with brain stimulation. Future testing of brain stimulation at a longer time point (perhaps 48 hours) from LSD would help to reduce interactions with acute effects; it is worth noting that the majority of the neuroplastic processes elicited by LSD that we are trying to interact with occur within 1 week.^64^

As work has shown enhanced synaptogenesis and dendritogenesis with similar doses of LSD as used here, we did not replicate the microscopic characterization of enhanced neuroplasticity. However, as we continue to characterize the interaction of psychedelic drugs and brain stimulation, it will be important to determine if adding brain stimulation alters the structural changes in neurons reported from LSD alone.

Although we balanced our groups to include roughly equal numbers of males and females, we did not have enough samples to adequately power comparisons between sexes. In humans, sex is not predictive of the effects or optimal dosing of psychedelic drugs including LSD.^65^ However, in rats there is some evidence that LSD and other psychedelic drugs have slightly different behavioral effects in males and females.^66, 67^ Thus, future work should continue to include both sexes and when possible be powered to compare between sexes directly.

We used a relatively high dose of LSD (0.15 mg/kg) in this experiment in attempts to maximize the potential interaction between enhanced neuroplasticity and brain stimulation. However, it is vital that a more complete dose-response relationship is teased out by also testing a series of other doses. This is important not only to determine the minimum effective dose, but also because the pharmacodynamics of LSD are partially determined by dose; for example, lower doses (<0.02 mg/kg) of LSD have less of an effect on dopaminergic neurons.^68^ Further, as LSD is a pharmacologically promiscuous drug it is unclear if 5HT-2A signaling is necessary or sufficient to produce the observed effects.^44^

We limited our data analysis of the acute effects of LSD to the last 30 minutes of recordings to account for the acute ataxia caused by LSD. While the amount of time spent resting was not significantly different between LSD and SAL rats in the final 30 minutes, there was still a considerable amount of variation in time spent resting across rats given LSD (**Supplemental Figure 1**) which could have contributed to the reported LFP differences. We did not score resting behavior beyond the acute recordings as even scoring behaviors at a fine level of resolution (separating out many different behaviors beyond just “resting”) would still fail to capture all possible behavioral and physiological differences between groups that could confound interpretation of LFP differences. We acknowledge that LSD+stim could be inducing different behavioral/physiologic changes compared to SAL+stim which is then reflected indirectly in recorded LFP differences. Importantly, we still interpret this to mean that there is a significant difference in what LSD+stim is doing to brain activity compared to SAL+stim, it just cannot be determined if the induced LFP changes are direct or indirect outcomes of stimulation. There is increasing interest in the discovery and testing of other compounds capable of inducing neural plasticity, especially compounds without the acute perceptual effects of psychedelic drugs.^69^ Beyond structural imaging of neurons to assess neuroplasticity, we hypothesize that pairing these new compounds with brain stimulation could provide a neural systems-level readout of latent plasticity.

## Conclusion

We have further confirmed the findings of acutely decreased low frequency power across the brain when rats are given LSD. We have extended these findings to show that despite these acute effects disappearing after 24 hours, there are still latent effects of LSD that interact with brain stimulation to create different changes in brain activity compared to brain stimulation alone. Combined with the knowledge that LSD opens a window of enhanced neural plasticity, our findings are the first to support the theory that psychedelic drugs could have a role clinically in combination with brain stimulation (e.g., TMS, electroconvulsive therapy, or deep brain stimulation) to achieve enhanced effects on brain activity and relevant clinical outcomes.

## Data Sharing

All processed LFP data and code necessary for analysis has been made available on Gnode (10.12751/g-node.xrxt9n).

## Author contributions

Lucas Dwiel: Conceptualization, data curation, formal analysis, funding acquisition, investigation, methodology, project administration, visualization, writing - original draft

Wilder Doucette: Conceptualization, funding acquisition, methodology, project administration, supervision, writing - review & editing

Angela Henricks: Funding acquisition, methodology, writing - review & editing Elise Bragg: Methodology, resources

---

**Supplemental Figure 1.**
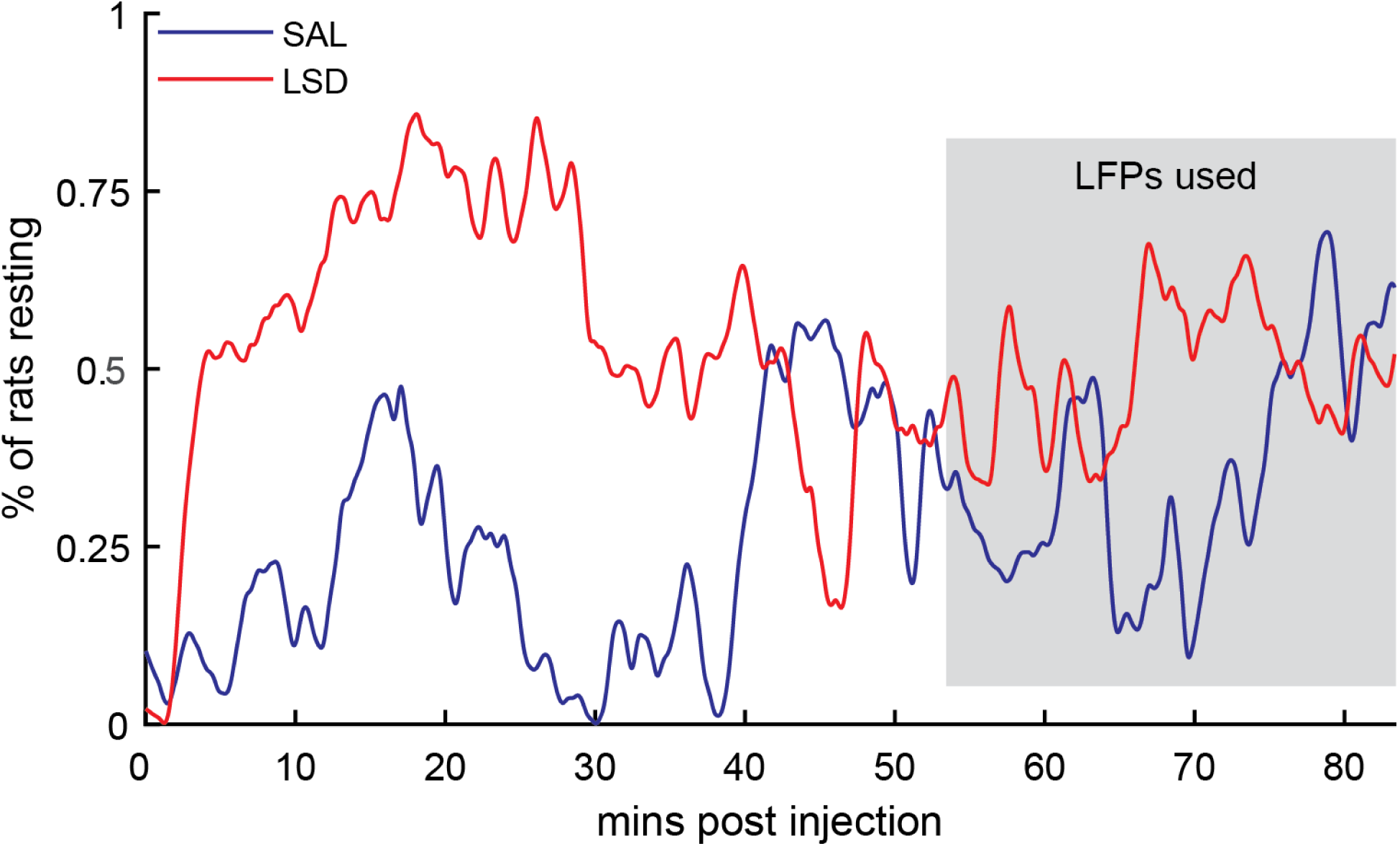
Percent of rats resting throughout recordings. Approximately 55 minutes after injections, the acute ataxia caused by LSD has worn off and the two groups of rats (LSD, red; SAL, blue) are spending a similar amount of time resting. LFPs from the last 30 minutes of the recording (grey box) were used to differentiate LSD and SAL.

**Supplemental Figure 2.**
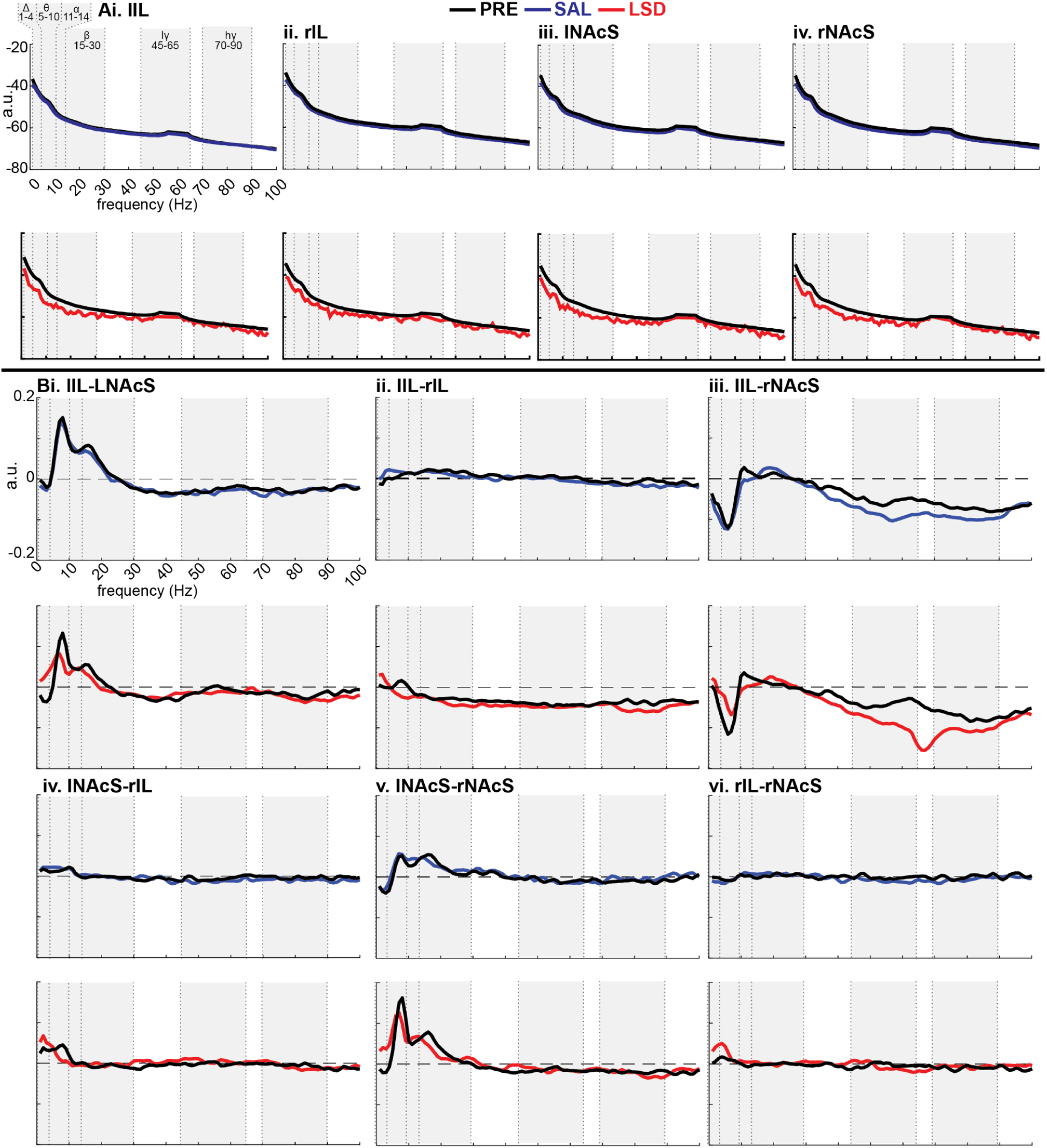
Power and coherence before and after saline (SAL) or LSD. **Ai-iv** Power from the four recording sites with data from before injections in black, after SAL in blue (top), and after LSD in red (bottom). Bi-vi Coherence from the 6 pairs of recording sites with data from before injections in black, after SAL in blue (top), and after LSD in red (bottom). N.B. the SAL and LSD data come from the end of the recording, ∼55 minutes after injections (see Supplemental Figure 1). Right, r; left, l; infralimbic cortex, IL; nucleus accumbens shell, NAcS; delta, Δ; theta, θ; alpha, α; beta, β; low gamma, lγ; high gamma, hγ.

**Supplemental Figure 3.**
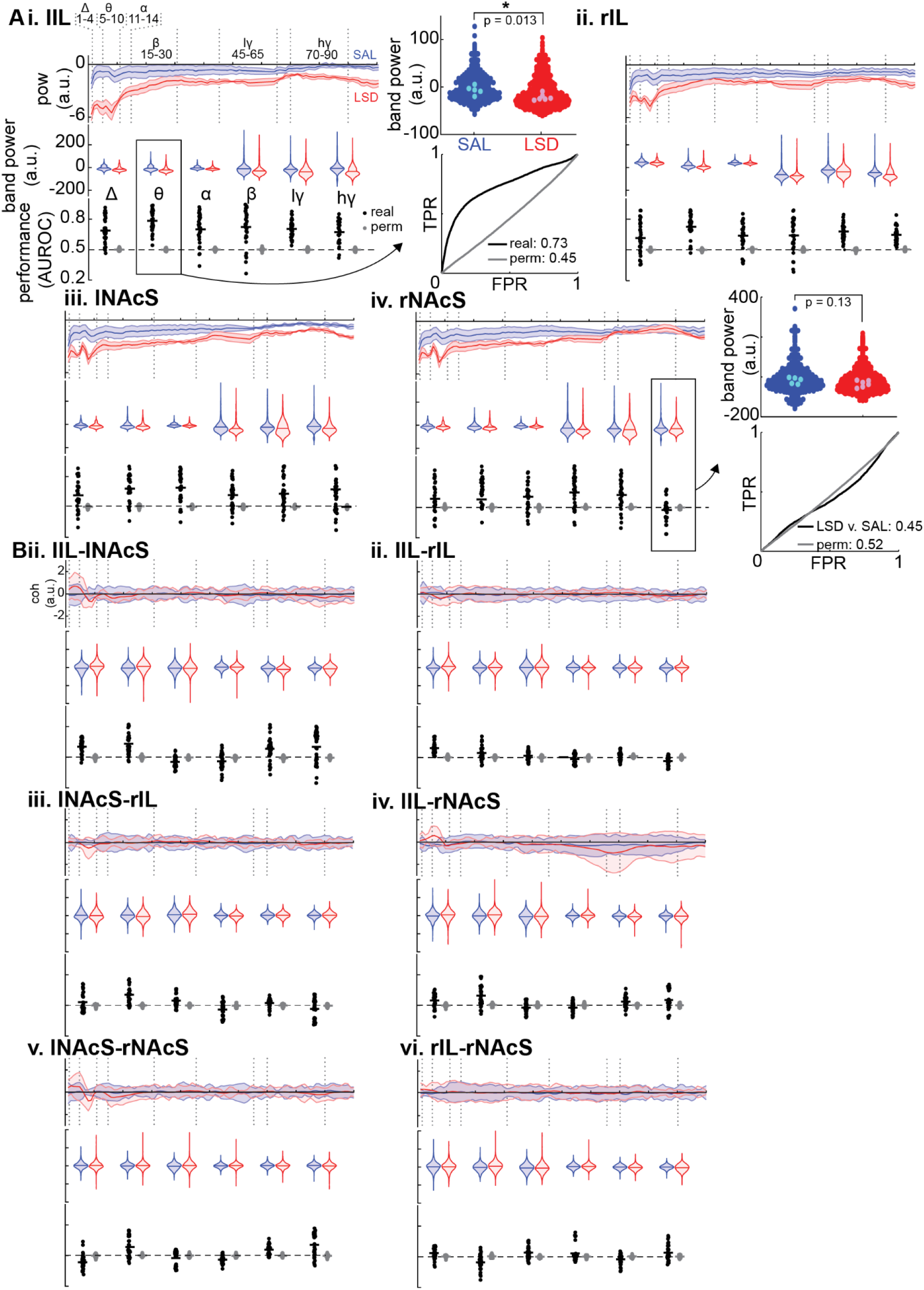
Power and coherence changes from pre- to post-injection of either saline (SAL) or LSD. **A** Top: change in power from pre to post injection of either SAL (blue) or LSD (red) in arbitrary units (a.u.) with ±1 standard deviation shaded. Middle: the difference in power between the SAL and LSD (black) with ±1 standard deviation shaded. Bottom: the distribution of single feature model AUCs from leave-one out testing (black) and permuted data (gray); horizontal lines indicate means. Brain regions are denoted as either right (r) or left (l) such that lIL (**Ai**) is the left infralimbic cortex. **B** Changes in coherence between SAL and LSD organized as in **A** with pairs of brain regions denoted such that lIL-lNAcS (**Bi**) is coherence between left infralimbic cortex and left nucleus accumbens shell.

**Supplemental Figure 4.**
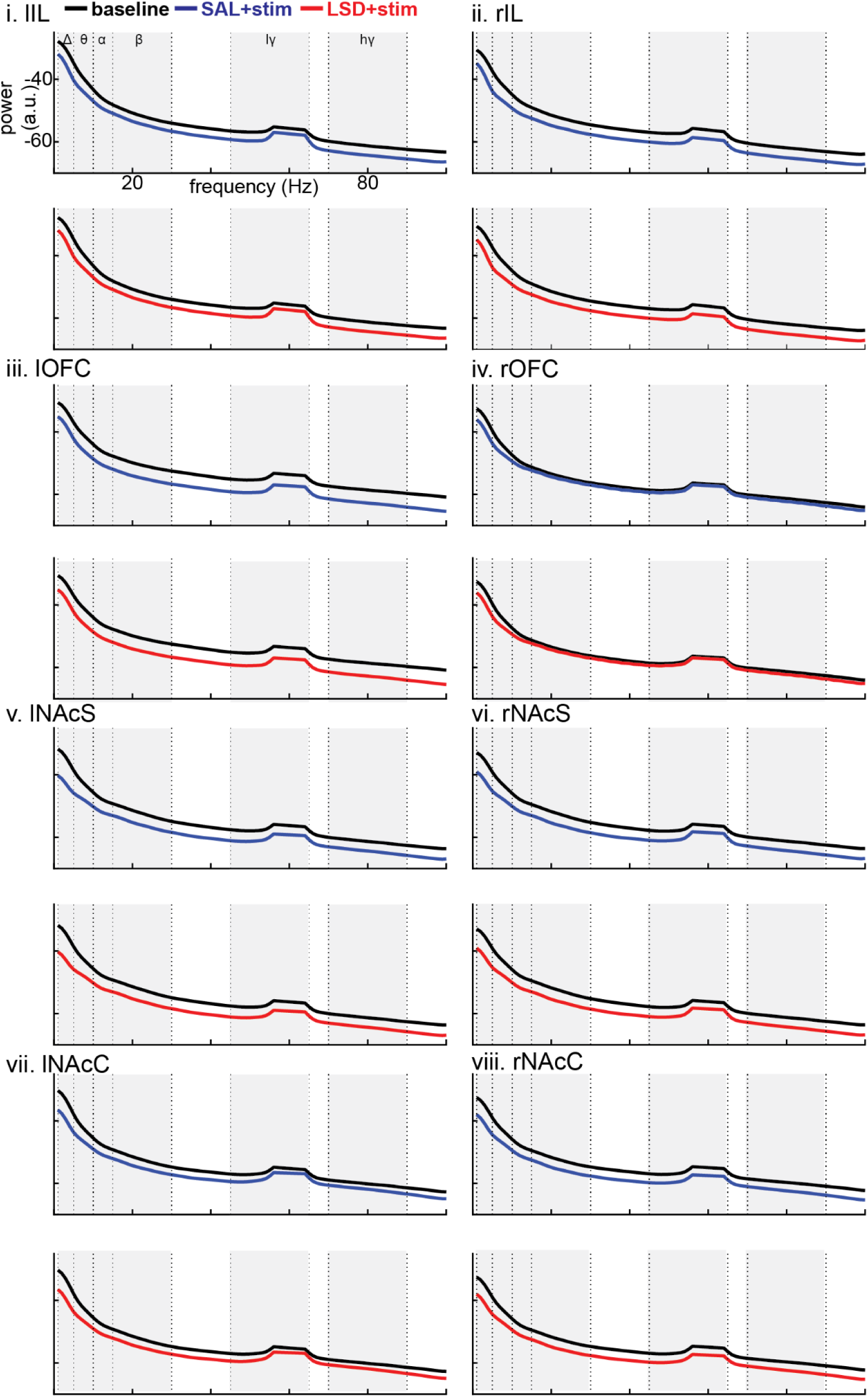
Power before and during saline+brain stimulation (SAL+stim) or LSD+stim. **Ai-iv** Power from the four recording sites with data from before injections in black, after SAL in blue (top), and after LSD in red (bottom). **Bi-vi** Coherence from the 6 pairs of recording sites with data from before injections in black, after SAL in blue (top), and after LSD in red (bottom). Right, r; left, l; infralimbic cortex, IL; nucleus accumbens shell, NAcS; delta, Δ; theta, θ; alpha, α; beta, β; low gamma, lγ; high gamma, hγ.

**Supplemental Figure 5.**
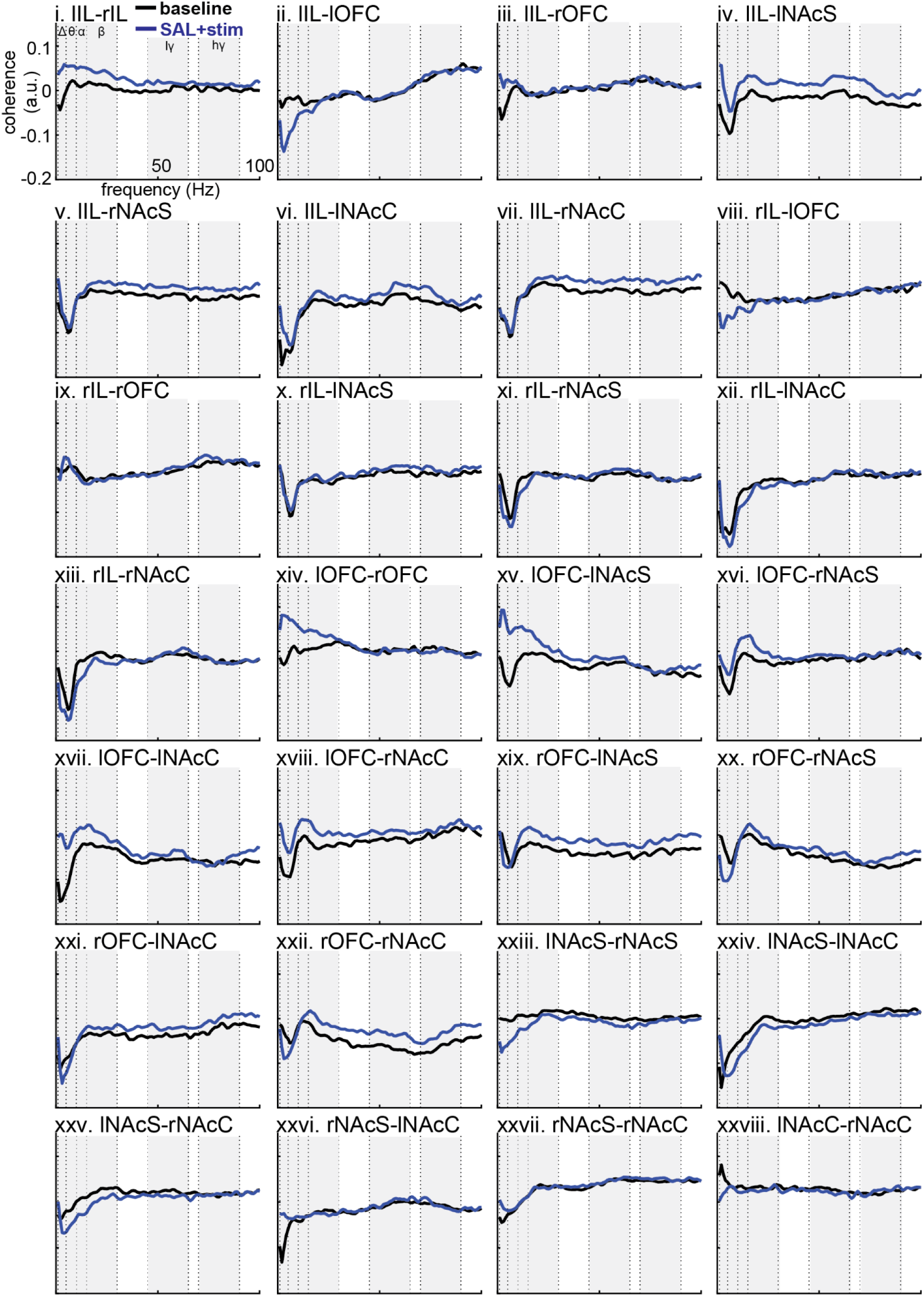
Coherence before and during saline+brain stimulation (SAL+stim). **i-xxviii** Coherence from the 28 recording site pairs with data from before stimulation in black (baseline), and after stimulation in blue (SAL+stim). Right, r; left, l; infralimbic cortex, IL; nucleus accumbens shell, NAcS; orbitofrontal cortex, OFC; nucleus accumbens core, NAcC; delta, Δ; theta, θ; alpha, α; beta, β; low gamma, lγ; high gamma, hγ.

**Supplemental Figure 6.**
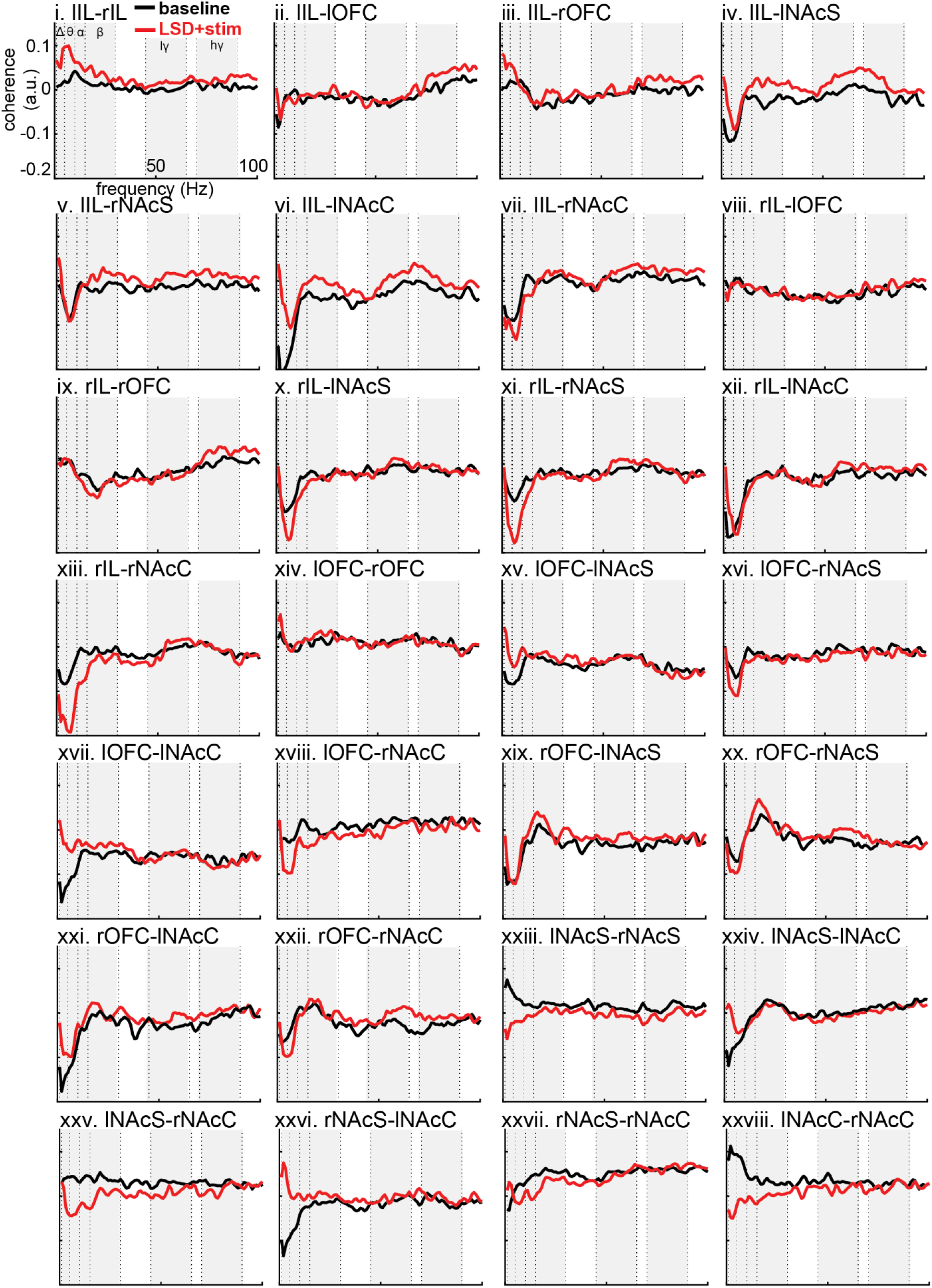
Coherence before and during LSD+brain stimulation (LSD+stim). **i-xxviii** Coherence from the 28 recording site pairs with data from before stimulation in black (baseline), and after stimulation in red (LSD+stim). Right, r; left, l; infralimbic cortex, IL; nucleus accumbens shell, NAcS; orbitofrontal cortex, OFC; nucleus accumbens core, NAcC; delta, Δ; theta, θ; alpha, α; beta, β; low gamma, lγ; high gamma, hγ.

**Supplemental Figure 7.**
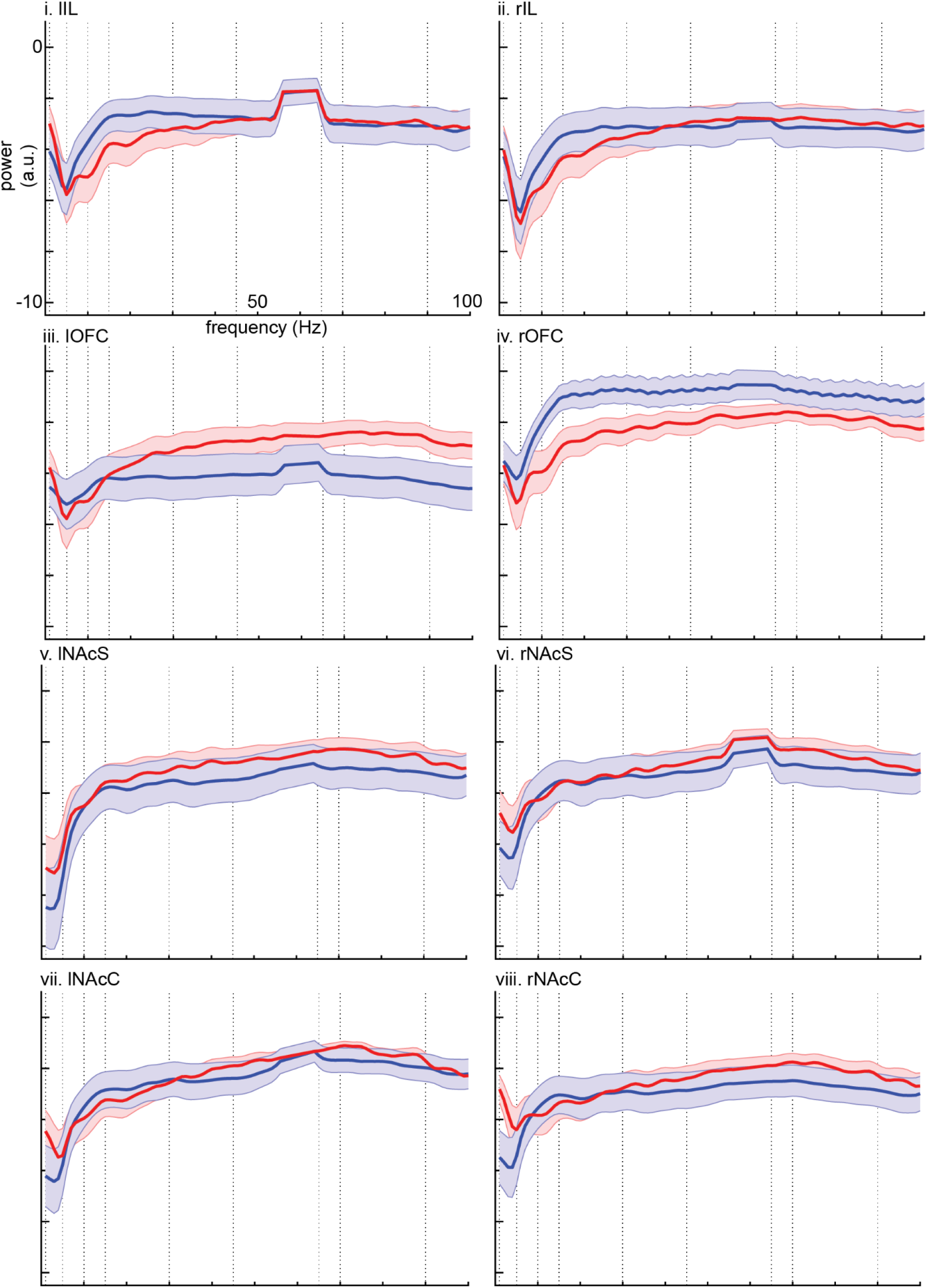
Power differences between before and during brain stimulation in rats given either LSD or SAL 24 hours prior. **i-viii** Power differences from the 8 recording sites with rats given SAL in blue (SAL+stim) and rats given LSD in red (LSD+stim). Right, r; left, l; infralimbic cortex, IL; nucleus accumbens shell, NAcS; orbitofrontal cortex, OFC; nucleus accumbens core, NAcC; delta, Δ; theta, θ; alpha, α; beta, β; low gamma, lγ; high gamma, hγ.

**Supplemental Figure 8.**
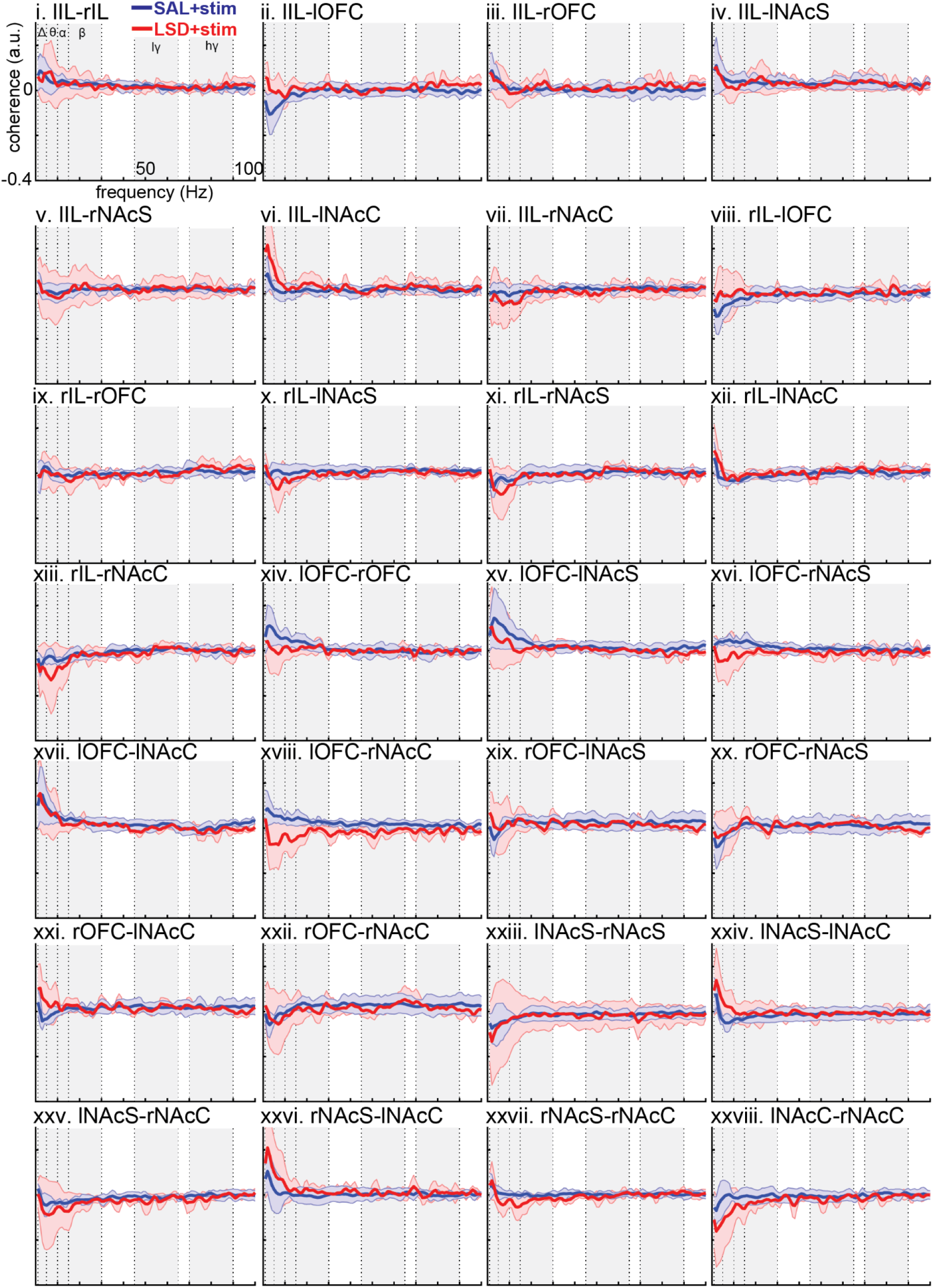
Coherence differences between before and during brain stimulation in rats given either LSD or SAL 24 hours prior. **i-xxviii** Coherence differences from the 28 recording site pairs with rats given SAL in blue (SAL+stim) and rats given LSD in red (LSD+stim). Right, r; left, l; infralimbic cortex, IL; nucleus accumbens shell, NAcS; orbitofrontal cortex, OFC; nucleus accumbens core, NAcC; delta, Δ; theta, θ; alpha, α; beta, β; low gamma, lγ; high gamma, hγ.

